# Concurrent measurement of perfusion parameters related to small blood and lymphatic vessels in the human brain using dynamic dual-spin-echo perfusion (DDSEP) MRI

**DOI:** 10.1101/2022.09.26.509366

**Authors:** Di Cao, Yuanqi Sun, Pan Su, Jay J. Pillai, Ye Qiao, Hanzhang Lu, Peter C.M. Van Zijl, Linda Knutsson, Jun Hua

## Abstract

**PURPOSE:** Accumulating evidence from recent studies has indicated the importance of studying the interaction between the microvascular and lymphatic systems in the brain. To date, most imaging methods can only measure blood or lymphatic vessels separately, such as dynamic-susceptibility-contrast (DSC) MRI for blood vessels and DSC MRI in the CSF (cDSC MRI) for lymphatic vessels. An approach that can measure both blood and lymphatic vessels in a single scan will offer the advantages such as halved scan time and contrast dosage. Based on previous works on DSC and cDSC MRI, this study proposes an MRI approach for concurrent measurement of perfusion parameters related to small blood and lymphatic vessels in the brain within one single scan.

**METHODS:** Bloch simulations were performed to optimize a dual-echo sequence for the measurement of gadolinium(Gd)-induced blood and CSF signal changes using a short and a long echo time, respectively. MRI experiments were performed in healthy subjects to evaluate the dual-echo approach by comparing it with existing separate methods.

**RESULTS:** The proposed method showed consistent results in human brains as previous studies using separate methods. Signal changes from small blood vessels occurred faster than lymphatic vessels after intravenous Gd-injection.

**CONCLUSION:** Gd-induced signal changes in blood and CSF can be detected simultaneously in healthy subjects with the proposed sequence. To the best of our knowledge, this may be the first study in which the temporal difference in Gd-induced signal changes from small blood and lymphatic vessels after intravenous Gd-injection was measured in the same human subjects.

## Introduction

The cerebrovascular system and cerebrospinal fluid (CSF) circulation system are two important components in the central nervous system (CNS). While the cerebrovascular system provides blood supply to the brain, CSF circulation is critical for waste clearance from the brain and immune surveillance (1). Recent studies in animals and humans have provided more details regarding CSF transportation and drainage in the CNS (1). First, the glymphatic system has been proposed, which aids CSF and interstitial fluid (ISF) transportation in brain parenchyma from the periarterial space to the perivenous space mainly through the aquaporin-4 (AQP4) membrane protein localized primarily in the end feet of astrocytes (2). Second, cerebral vessels with typical endothelial markers found in lymphatic vessels in other organs have been identified in several parts of the brain. These include the dura matter alongside the dural venous sinuses, regions around the middle meningeal artery and cribriform plate (3-5), areas in the basal part of the skull (6), and several other brain regions in animal models. Some of the meningeal lymphatic vessels have also been visualized in human brain (5). These cerebral lymphatic vessels may communicate with the glymphatic system and other routes for CSF circulation and are believed to play a crucial role in the drainage of CSF and ISF from brain tissues to cervical lymph nodes (3-9). The study of CSF flow in cerebral lymphatic vessels may provide essential information regarding the clearance of abnormal proteins and metabolites from brain tissues. The methodology developed in the current study is focused on cerebral lymphatic vessels, not the glymphatic system.

Accumulating evidence has indicated the interaction between the microvascular and lymphatic systems in the brain. As the CSF and ISF space in brain parenchyma lie alongside small blood vessels, CSF and ISF in the perivascular space are believed to be primarily driven by the arterial pulsation wave from these small blood vessels (7-9). A change in blood flow may lead to impaired lymphatic clearance, both of which may contribute to the pathogenesis of various brain diseases. Therefore, it is of importance to investigate the relationship between the microvascular and lymphatic systems in the brain using *in vivo* imaging methods that can capture the two systems in the same subject. To date, most imaging methods can only measure blood or lymphatic vessels separately. An approach that can measure the two systems in a single scan will offer the advantage of significantly shortened scan time, and more importantly, less confounding effects from physiological variations between scans. For MRI methods based on contrast agents, a combined method will reduce the number of doses of contrast media that need to be administrated in the participants.

Dynamic susceptibility contrast (DSC) MRI is a standard perfusion technique performed routinely in clinical MRI exams (10,11). It can measure several key parameters reflecting the function and integrity of blood vessels, such as cerebral blood flow (CBF) and volume (CBV), and blood brain barrier (BBB) permeability. In typical DSC MRI experiments, MR images are continuously acquired before, during and after intravenous (IV) administration of Gadolinium (Gd) based contrast agents, generating a response curve from which the microvascular parameters can be derived. Gd-based contrast agents are impermeable to an intact BBB in cortical blood vessels. Nevertheless, as the dural blood vessels lack such a BBB, the Gd-based contrast agents can cross the dural blood vessel wall and enter the CSF. Therefore, post-contrast MR signal changes can often be observed in the CSF at certain locations within the intra-cranial space (12-15). To date, DSC MRI has been optimized primarily for detecting Gd-induced contrast changes in blood vessels (10). Recently, we developed an MRI approach (16) for the detection of Gd-based signal changes in the CSF, namely DSC MRI in the CSF (cDSC MRI), which can be used to measure perfusion parameters related to cerebral lymphatic vessels in the human brain. Most existing approaches for the measurement of Gd-induced MR signal changes in the CSF and cerebral lymphatic vessels take at least a few minutes to achieve whole brain coverage and sufficient spatial resolution, which provides a relatively low temporal resolution (5,8,17-26). The cDSC MRI method was optimized to detect dynamic signal changes in the CSF and cerebral lymphatic vessels with a sub-millimeter spatial resolution, a temporal resolution of less than 10 seconds, and whole brain coverage (16).

In the current study, we propose a new MRI approach for concurrent measurement of perfusion parameters related to small blood and lymphatic vessels in the brain within one single scan, termed “dynamic dual-spin-echo perfusion (DDSEP) MRI”. A dual-echo turbo-spin-echo (TSE, also known as fast spin echo or FSE) sequence was optimized for the measurement of Gd-induced blood and CSF signal changes using a short and a long echo time (TE), respectively. In this first proof-of-concept study, the proposed DDSEP MRI approach was implemented with a multi-slice version, in which 3 slices covering some of the brain regions where cerebral lymphatic vessels have been identified were acquired. Theoretical simulations were performed to optimize the imaging parameters. MRI experiments were performed in healthy human subjects to evaluate the proposed combined approach by comparing it with existing separate methods. Approaches to expand the current sequence to a whole brain scan are discussed based on the results obtained from the current study.

## Methods

### Pulse sequence

In order to detect blood and CSF signals in a single MRI scan, MR signals from blood and CSF need to be separated based on their substantially different T1 and T2 values (**Table 1**). In addition, the spatial and temporal resolutions need to be optimized to balance between the two types of measures. Specifically, the sequence design needs to fulfill the following criteria:

**Table 1.**
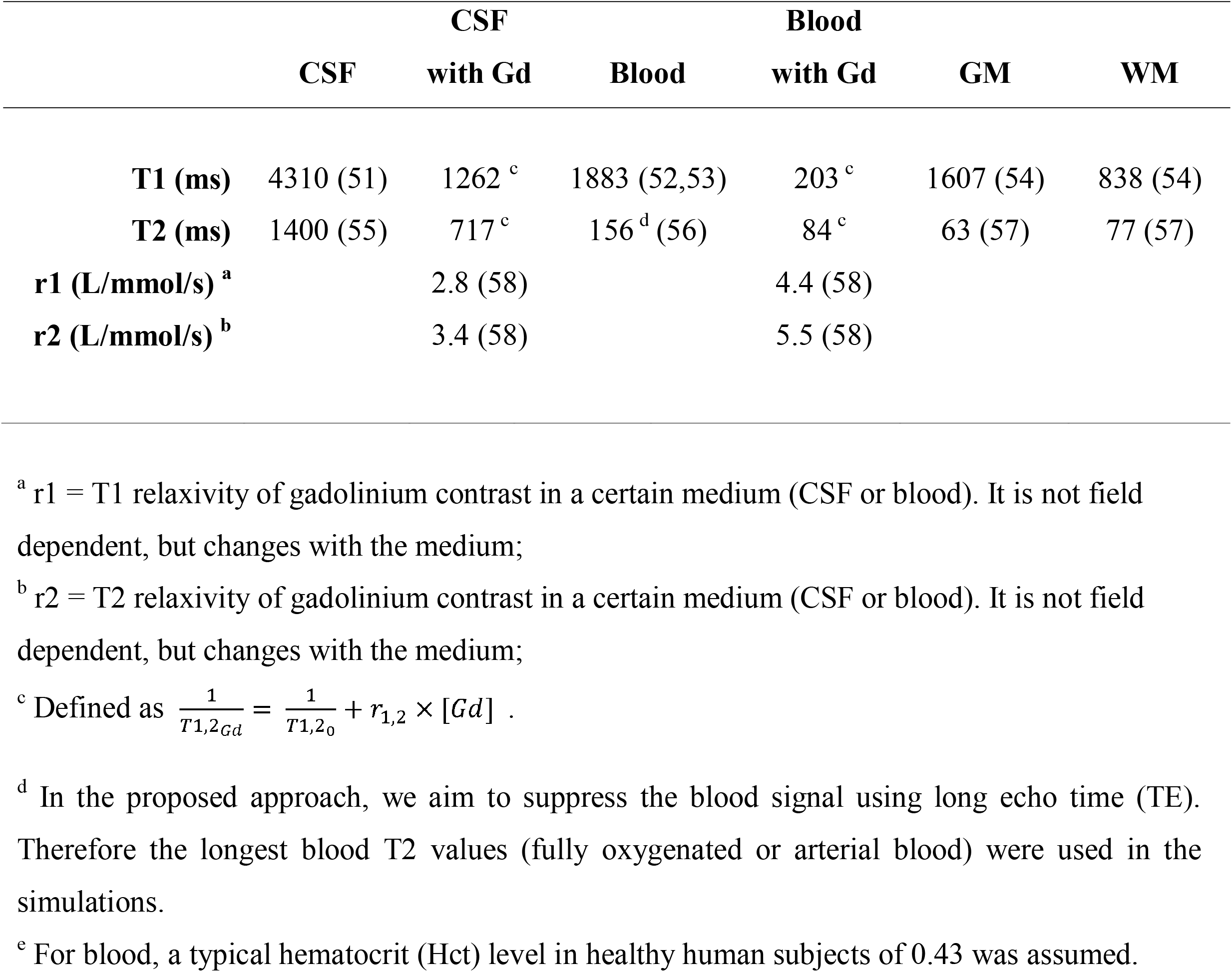
Literature values for key parameters used in the simulations for 3T.

1. For blood vessels, a relatively high temporal resolution of 1.5–2.5 s is needed to capture the dynamic changes. A spatial resolution of 2-3 mm is sufficient as in typical DSC MRI scans for measuring blood perfusion parameters. A relatively short TE (~80 ms for spin echo) is considered optimal for blood signals.
2. For CSF in cerebral lymphatic vessels, a relatively low temporal resolution of 5–10s is acceptable, based on the typical onset time measured in previous studies (16). A sub-millimeter spatial resolution is ideal to minimize partial volume effects as cerebral lymphatic vessels often run alongside small blood vessels, although the voxel size does not need to be smaller than the microvessels if the signals from parenchyma and blood can be effectively suppressed (~0). As CSF T2 is much longer than that of parenchyma and blood in the brain (**Table 1**), a long TE (~ 500 ms) is required to minimize signals from parenchyma and blood.

It should provide sufficient MR signal contrast in blood and CSF before and after Gd administration.

According to these criteria, we propose a dual-echo TSE sequence for simultaneous measurement of blood and CSF signal changes in the brain (**Figure 1**). Similar methods have been used in DSC MRI for measuring dynamic signal changes from blood vessels (10). In our proposed approach, the following unique features were implemented based on the requirements described above:

**Figure 1.**
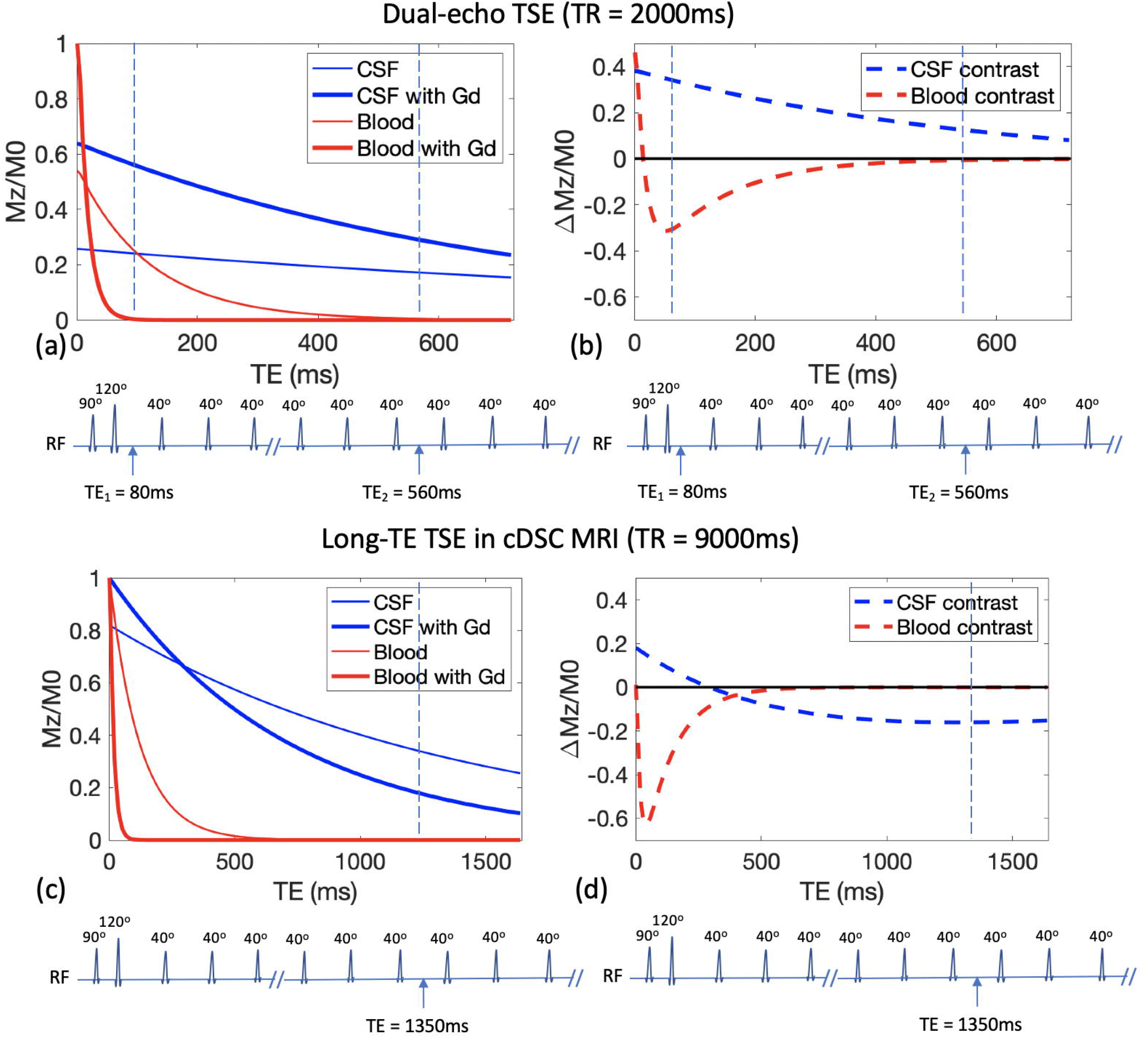
Pulse sequences and simulation results for the proposed dual-echo TSE sequence **(a, b)**, and the long-TE TSE sequence developed in the previous cDSC MRI study **(c**,**d)**. MR signals in blood and CSF with and without Gd are simulated in **(a, c)** and the contrast between pre- and post-Gd signals in blood and CSF are shown in **(b, d)**. The fractional MR signals (Mz/M0) and the fractional MR signal changes before and after Gd injection (ΔMz/M0=(Mz_post_Gd-Mz_pre_Gd)/M0) are displayed as functions of TE. The vertical dashed lines indicate the TEs in respective sequences.

1. T1-dominant contrast in the CSF: Gd shortens both T1 and T2 values in blood and CSF, which affects the MR signals in the proposed sequence oppositely. In our previous work on cDSC MRI (16), two types of sequences were evaluated for detecting Gd-induced MR signal changes in the CSF. When a relatively short repetition time (TR) is used, the TSE sequence furnishes a T1-dominant positive contrast in CSF after Gd injection. When a relatively long TR is used, the TSE sequence shows a T2-dominant negative contrast in CSF after Gd injection, which was employed in cDSC MRI (16). However, in the proposed combined sequence here, a T1-dominant sequence with short TR was chosen for measuring Gd-induced MR signal changes in both blood and CSF mainly due to the temporal resolution (TR) requirement (see the next point).
2. A temporal resolution of 2 s and a spatial resolution of 1×1×2 mm^3^ were chosen to balance the requirements between blood and CSF signals. The temporal resolution is typical for measuring dynamic changes in blood vessels in DSC MRI and is sufficient for detecting dynamic changes in cerebral lymphatic vessels in human brains based on previous data (16). This is the primary reason for choosing the T1-dominant sequence with short TR in the combined method. Thus, we have a T1-dominant contrast in CSF and a T2-dominant contrast in blood, which has much shorter T1 values than CSF. Note that the spatial resolution here is much finer than the typical spatial resolution of ~2×2×5mm^3^ in DSC MRI.
3. A short and a long TE were used in the proposed dual-echo TSE sequence. The first echo (short TE1 = 80 ms) is intended to measure the blood signals similar to DSC MRI. The second echo (long TE2 = 560 ms) measures predominantly the CSF signals with parenchyma and blood signals suppressed.
4. Theoretical simulations (please see the next section) were performed to optimize the imaging parameters in the proposed sequence in order to maximize MR signal contrast in blood and CSF before and after Gd administration.
5. The TSE sequence used is a spin-echo (SE) based sequence, which has better spatial specificity for Gd-induced signal changes (27-30) compared to the commonly used gradient echo (GRE) based sequences. This is important as cerebral blood and lymphatic vessels are often adjacent to each other (3-5).

### Simulations

Bloch simulations were performed to calculate the MR signal and contrast before and after Gd injection from the proposed pulse sequence in blood and CSF. In-house code programmed in Matlab (MathWorks, Natick, MA, USA) was used for the simulations. **Table 1** summarizes the literature values for the parameters used in the simulations. The relaxation times of CSF after Gd injection were calculated based on the relaxivity values r1 and r2 (**Table 1**) for the Gd contrast medium Prohance (or Gadoteridol; Bracco S.p.A., Milan, Italy; 0.5 mmol/ml) used in this study with a standard dosage of 0.1 mmol/kg. The concentration of Gd in blood was assumed to be 10 mmol/L for the bolus phase typically measured in DSC MRI (10). The concentration of Gd in CSF was assumed to be 0.2 mmol/L, based on the results in healthy human subjects from the literature (16). A second simulation was performed with varying Gd concentrations in blood and CSF to investigate its effects on the MR signals.

### Quantification of Gd concentration ([Gd]) in the CSF

Since the quantification of Gd concentration in the blood has been extensively studied in DSC and dynamic contrast enhancement (DCE) MRI, the current study is focused on Gd concentration in the CSF. In the original cDSC MRI, the CSF contrast is T2-dominant and decreases monotonically with Gd concentration, which makes it possible to determine Gd concentration from the CSF contrast (16). However, in DDSEP MRI, the CSF contrast becomes T1-dominant due to the requirement for a short TR, which shows a biphasic relationship with Gd concentration (please see simulation results). Thus, Gd concentration cannot be uniquely determined from the long-TE CSF images alone. The CSF signal changes measured at the short TE in DDSEP MRI increase with [Gd] monotonically (please see simulation results), when the Gd concentration is within the range expected for the CSF and cerebral lymphatic vessels in the human brain after intravenous (IV) Gd administration with a standard dose. But the short-TE images in DDSEP MRI have signal contributions from both blood and CSF. Based on our data measured in healthy human subjects in the current study (please see human subject results), the blood and CSF signal changes at the short TE showed significant temporal separation for at least 20 seconds. When the CSF signals showed significant Gd-induced changes, the blood signals have returned to its pre-Gd level. Based on these assumptions, Gd concentration [Gd] for the time period of 30 seconds after Gd injection can be uniquely solved from (S1_Gd_-S1_0_)/S2_0_, where S1_0_ and S1_Gd_ represent MR signals measured at the short TE before and after Gd injection, respectively; and S2_0_ represents the long-TE signal measured before Gd injection. Note that S2_0_ can be obtained from the long TE in DDSEP MRI, or it can simply be acquired using a separate scan with the same parameters before Gd injection. The complete derivation is shown in the **Supporting Information**.

### MRI and Gd contrast

MRI scans were performed on a 3 Tesla (3T) Philips human MRI scanner (Philips Healthcare, Best, The Netherlands). A 32-channel phased-array head coil was used for signal reception and a dual-channel body coil for transmit. The Gd contrast agent (Prohance, or Gadoteridol; Bracco S.p.A., Milan, Italy) was administered intravenously (IV) using a standard procedure (dosage=0.1mmol/kg, injection rate=5mL/s).

### Human studies

Eight healthy volunteers were recruited for this study (three female and five male, age 23-36 years, 27±6 years). All participants were screened with a creatinine measurement of less than 1.4 mg/dl on whole blood to be eligible for an MRI scan with Gd. This study was approved by the Johns Hopkins Institutional Review Board, and written informed consent was obtained from each participant. The parameters for the proposed dual-echo TSE scan were chosen based on the simulation results.

The following scans were performed on 3T for each subject sequentially: **1)** a 3D fluid-attenuation inversion recovery (FLAIR) sequence similar to previous cerebral lymphatic vessel studies (5): TR/TI/TE = 4800/1650/307 ms, TSE factor =167, Compressed-Sensing SENSE (CS-SENSE) = 8.5, voxel = 1mm isotropic, 130 slices, scan duration = 2 minutes and 10 seconds; **2)** a 2D multi-slice FLAIR sequence similar to previous cerebral lymphatic vessel studies (5) but with the same spatial resolution and coverage as the proposed dual-echo TSE sequence in this study: TR/TI/TE = 4800/1650/200ms, TSE factor =41, CS-SENSE = 2, voxel = 1×1×2mm^3^, 3 slices, scan duration = 48 seconds; **3)** the proposed dynamic dual-echo TSE sequence (the following parameters were chosen based on the simulation results): TR/TE1/TE2/echo spacing (ES) = 2000/80/560/5.4ms, TSE factor = 120, number of shots = 1, partial Fourier = 0.7, CS-SENSE = 2, voxel = 1×1×2 mm^3^, 3 slices, and the sequence was performed continuously while 150 volumes of images were acquired immediately before and after Gd injection, respectively, total scan duration = 10 minutes, head specific absorption rate (SAR) < 15% of the FDA limit; **4)** a second 2D multi-slice FLAIR scan (post-Gd injection) with parameters identical to the first one.

Data acquired using the proposed sequence were compared with data from existing separate sequences: cDSC MRI for lymphatic vessels and standard DSC MRI for blood vessels. Because normally only one standard dose of Gd contrast media is given to a participant during each MRI session, it is not possible to perform all three scans in the same subjects. Therefore, in this study, with the purpose of comparing the three techniques in healthy human subjects, existing data were used for comparison. For the cDSC scan, data from our previous study (16) in eight healthy human subjects (four female and four male, age 23-63 years, 43±17 years) were used. Briefly, for reference, this dynamic 3D TSE cDSC MRI scan was performed with the following parameters: TR/TE/ES = 9000/1289/4.4ms, TSE factor = 950, number of shots = 1, partial Fourier = 0.75×0.7, CS-SENSE = 12, voxel = 0.75mm isotropic, 173 slices, and the sequence was performed continuously while 33 and 107 volumes of whole brain images were acquired immediately before and after Gd injection, respectively, total scan duration = 21 minutes. In addition, a 3D FLAIR scan was performed before and after Gd injection with the following parameters: TR/TI/TE = 4800/1650/307ms, TSE factor =167, CS-SENSE = 8.5, voxel = 1mm isotropic, 130 slices, scan duration = 2 minutes and 10 seconds. For standard DSC MRI, data from our previous study (31) in eight healthy human subjects (four female and four male, 26 ± 4 years old,) were used. The DSC MRI scan was performed using the following parameters: 2D GRE echo-planar-imaging (EPI), TR/TE = 1000ms/30ms, flip angle = 54°, voxel = 2.8×2.8×5 mm^3^, slice gap = 0.5 mm, 18 slices, scan duration = 72 seconds. In addition, a T1-weighted magnetization-prepared rapid acquisition gradient-echo (MPRAGE) was performed with the following parameters: TR/TE/TI = 8.1/3.7/1100 ms, shot interval = 2100 ms, flip angle = 12°, voxel = 1×1×1 mm^3^, 160 slices, sagittal slice orientation, and scan duration = 3 minutes and 57 seconds.

Note that the durations of the DDSEP and cDSC scans were chosen to be substantially longer (10-20 minutes) than needed for investigational purposes. Based on the time courses measured in our previous study (16) and the current study, for future clinical applications, the same scans only need to be performed for about 5 minutes to capture the dynamic signal changes before and after Gd administration.

### Data analysis

The statistical parametric mapping (SPM) software package (Version 12, Wellcome Trust Centre for Neuroimaging, London, United Kingdom; http://www.fil.ion.ucl.ac.uk/spm/) and other in-house code programmed in Matlab (MathWorks, Natick, MA, USA) were used for image analysis. All dynamic images acquired before and after Gd injection were motion corrected using the realignment routine in SPM. FLAIR and MPRAGE images were co-registered to the corresponding mean dynamic images after realignment. Masks of cortical grey matter (GM) and white matter (WM) were obtained using the segmentation routine in SPM. Regions of interest (ROI) that cover the cerebral lymphatic vessels were manually delineated on the anatomical (FLAIR or MPRAGE) images for each subject, which were then overlaid onto all MRI scans from the same subject for further analysis. Similar to the data from the proposed DDSEP sequence, the ROIs of cDSC MRI and GRE EPI DSC MRI data acquired in different cohorts of healthy human subjects were identified in the individual space for each participant before subsequent analysis was performed.

The following parameters were computed for all dynamic imaging data. The relative signal change (ΔS/S) was calculated as the difference signal between pre- and post-Gd periods divided by average pre-Gd signal. The pre- and post-Gd signals were averaged over 52-250s before and 100-298s after Gd injection, respectively in cDSC and DDSEP MRI; and 4-12s before and 18-22s after Gd injection, respectively in DSC MRI. Temporal signal-to-noise ratio (tSNR) was taken as the signal divided by standard deviation along the time course in each voxel. Contrast-to-noise ratio (CNR) was defined as the product of tSNR and ΔS/S. A threshold of CNR>1 was applied in all dynamic scans to exclude voxels with high noise levels.

The short-TE images in the proposed approach and the standard DSC MRI scans were further analyzed using the Dynamic Susceptibility Contrast MR ANalysis (DSCoMAN) software (https://sites.duke.edu/dblab/dscoman/), from which typical cerebrovascular perfusion parameters such as CBV, CBF, mean transit time (MTT), and time to peak (TTP) were calculated in the entire grey matter region. In addition, in-house Matlab code was used to calculate ΔS/S, onset time (T_onset_), and return time (T_return_).

The long-TE images in the proposed approach and cDSC scans were further analyzed using in-house Matlab code, from which ΔS/S, onset time (T_onset_), and TTP were computed using the same procedure as in our previous study (16). Briefly, onset time (T_onset_) was defined as the time when significant signal change is detected in the dynamic TSE scans after Gd injection.

TTP was defined as the time when maximum signal change is reached in the dynamic TSE scans after Gd injection. Gd concentration ([Gd]) in the CSF after TTP (i.e. peak Gd concentration) was computed as outlined above and in the **Supporting Information**. Two ROIs where cerebral lymphatic vessels have been identified in human brains were chosen, including regions around the dural sinuses (superior sagittal sinus) and cribriform plate (3-5). The ROI around the dural sinuses covers the previously reported cerebral lymphatic vessels running alongside the superior sagittal sinus (3-5), which can be appreciated as vessels lateral to the superior sagittal sinus between the periosteal dura mater and meningeal dura mater in a coronal view on FLAIR images. The ROI around the cribriform plate covered the area right above the olfactory bulb and olfactory tract identified on FLAIR images. The ROI selection was performed by two experienced investigators (D.C. and J.J.P.).

Two-sample two-tailed t-tests with unequal variances were used for statistical analysis.

## Results

### Simulations

**Figures 1(a,b)** show the main simulation results for the proposed dual-echo TSE sequence. Fractional MR signals (Mz/M0) and the fractional MR signal changes before and after Gd injection (ΔMz/M0=(Mz_post_Gd-Mz_pre_Gd)/M0) were plotted as functions of TE. In the proposed approach, the pre-versus post-Gd blood contrast ΔMz/M0 (**Figure 1b**) becomes more negative with increasing TE at first, and then gradually returns to zero at longer TEs. The short TE in the proposed approach was chosen around the maximum pre-versus post-Gd blood contrast ΔMz/M0. The pre-versus post-Gd CSF contrast ΔMz/M0 (**Figure 1b**) shows a positive signal change in this T1-weighted sequence with a TR of 2s, and gradually decreases with increasing TE. The long TE in the proposed approach was chosen when the blood signals (**Figure 1a**) are completely suppressed (~0) in order to minimize the partial volume effects from small blood vessels that often run alongside cerebral lymphatic vessels. Note that at the short TE around the maximum pre-versus post-Gd blood contrast, positive CSF contrast can also be detected, the magnitude of which is comparable to that of the negative blood contrast at short TE. In comparison, **Figures 1(c,d)** show the simulation results for the original cDSC MRI approach developed in our previous study (16). Here, the pre-versus post-Gd blood contrast ΔMz/M0 (**Figure 1d**) shows a negative signal change that peaks around TE = 70 ms, but quickly returns to zero at longer TEs (TE > 600 ms). As this sequence was optimized for T2-weighted pre-versus post-CSF contrast ΔMz/M0 (16), a relatively long TR of 9s was chosen, which is too long for measuring blood perfusion. The pre-versus post-Gd CSF contrast ΔMz/M0 (**Figure 1d**) shows a negative signal change for TE > 500ms in this T2-weighted sequence (16) and peaks around TE = 1350 ms. The TE in cDSC MRI was chosen around the time when the CSF contrast peaks and the blood signals (**Figure 1c**) are negligible. The magnitude of the CSF contrast (~0.15) in cDSC MRI is smaller than that in the proposed DDSEP sequence at the long TE (~0.2, **Figure 1b**).

In all simulations in **Figure 1**, the concentration of Gd in blood and CSF was fixed based on the literature in healthy human subjects (see Methods). **Figure 2** shows simulation results for the pre-versus post-Gd contrast ΔMz/M0 in blood and CSF as a function of Gd concentration for both sequences. For the blood contrast (**Figure 2a**), the first echo (TE1) in the proposed DDSEP sequence shows a biphasic relationship with Gd concentration as the contrast is T1-dominant for blood signals. The blood contrast is zero for the second echo (TE2) in DDSEP and the original cDSC MRI as the TE is sufficiently long to suppress the blood signals. In the DDSEP sequence, a theoretical TE = 0 signal can be fitted numerically from the two echoes. At TE = 0, the contrast is only affected by the Gd induced T1 change, which leads to a positive signal change that quickly plateaus with increasing Gd concentration as the post-Gd blood signal approaches zero due to fast T1 decay (thus maximum pre-versus post-Gd contrast). For the CSF contrast (**Figure 2b**), both the first (TE1) and second (TE2) echo in the proposed DDSEP sequence show a biphasic relationship with Gd concentration as the contrast is T1-dominant for the CSF signals in this sequence with a short TR. By contrast, in the original cDSC MRI sequence, the T2-dominant contrast for the CSF signals (16) is always negative and increases in magnitude (more negative) monotonically with Gd concentration, and plateaus for approximately [Gd] > 0.5 mmol/L. At TE = 0 in the DDSEP sequence, similar to the blood contrast, the CSF contrast is also only affected by the Gd induced T1 change, which leads to a positive signal change that quickly plateaus with increasing Gd concentration. It is worth noting that for the typical range of Gd concentration in the CSF (0 - 0.5 mmol/L, shaded in **Figure 2b**) as measured in previous studies (16), only the second echo (TE2) in the proposed DDSEP sequence shows a biphasic relationship between the CSF contrast and Gd concentration, while for all other sequences, the CSF contrast either increases or decreases monotonically with Gd concentration, which makes it possible to determine Gd concentration from the signal changes.

**Figure 2.**
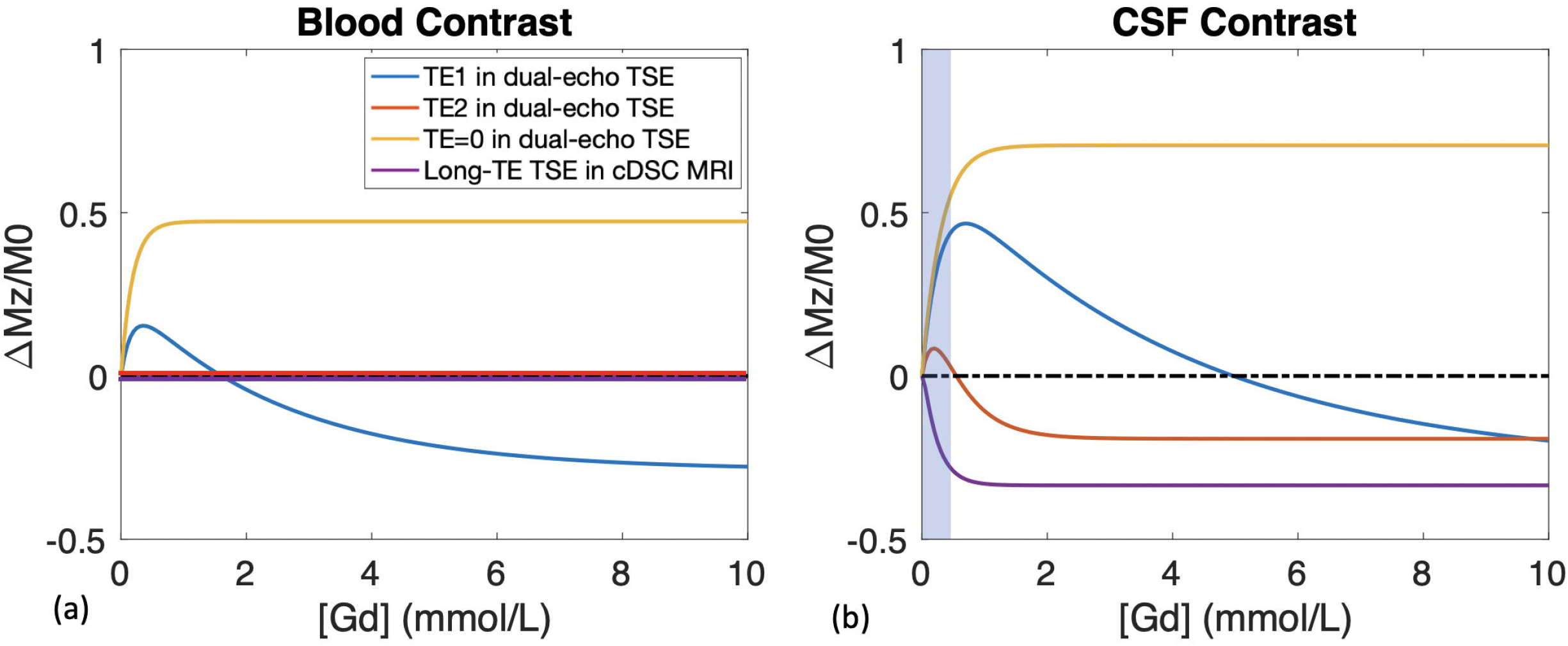
Simulations of the signal contrast with and without Gd (ΔMz/M0=(Mz_post_Gd-Mz_pre_Gd)/M0) in **(a)** blood and **(b)** CSF as a function of Gd Concentration ([Gd]) are demonstrated. The imaging parameters adopted in the human experiments are used for this simulation (see Methods). In the proposed dual-echo TSE sequence, signal contrasts at the two acquired TEs (TE1 = 80 ms, TE2 = 560 ms) and the theoretical TE = 0 (which can be numerically fit from TE1 and TE2) are calculated. Signal contrasts in the long-TE TSE in cDSC MRI (TE = 1350 ms) are also simulated.

### Human subject results: perfusion parameters related to blood vessels

**Figures 3-6** show typical images and time courses acquired in healthy human subjects on 3T. The quantitative results are compared in **Table 2**.

**Figure 3.**
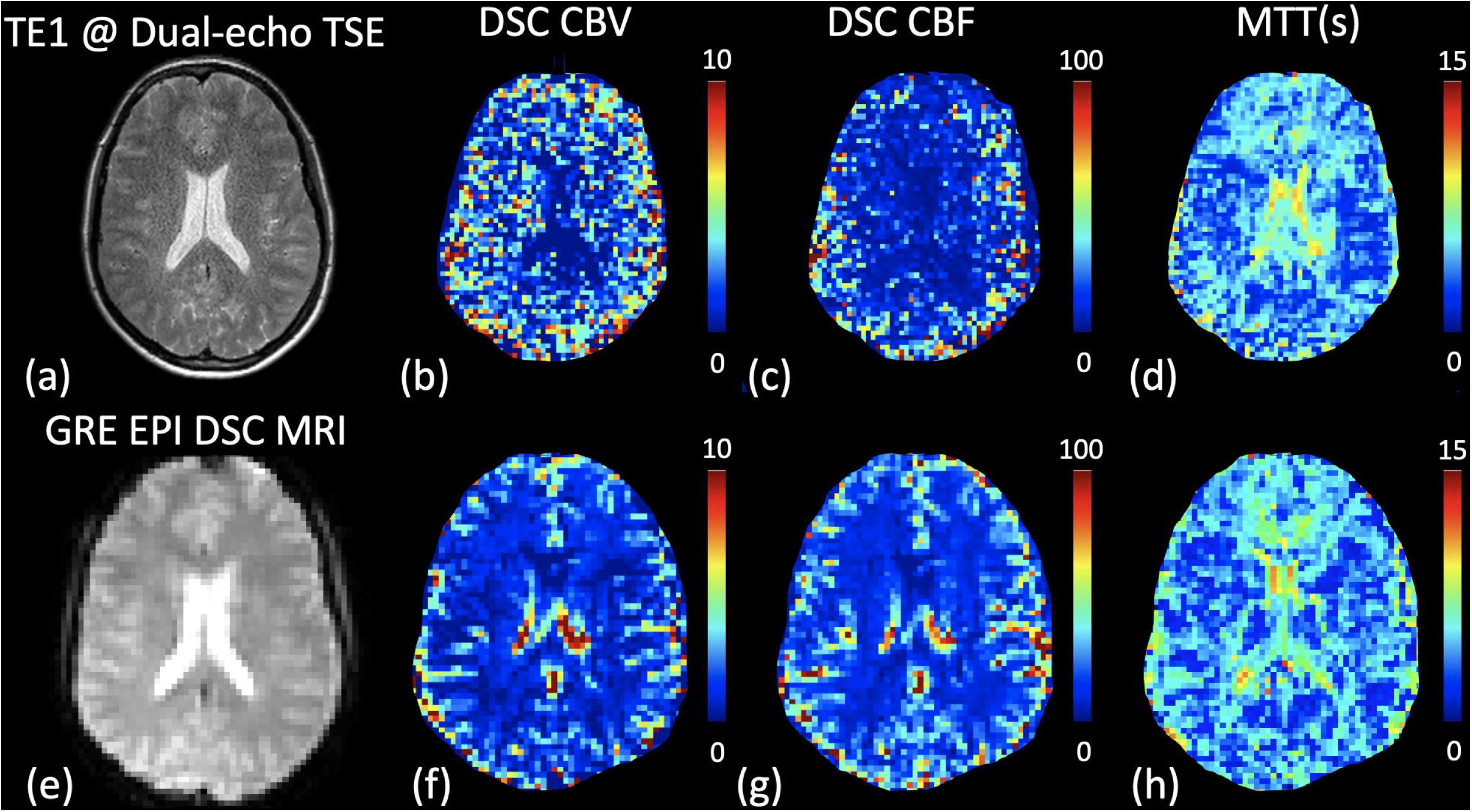
Representative results for the measurement of blood perfusion from human scans using the optimized dual-echo TSE sequence **(a-d)** and the standard GRE EPI DSC MRI sequence **(e-h)** on 3T are shown. The raw MR images from the short TE (TE1 = 80 ms) in the proposed dual-echo TSE sequence and from the standard GRE EPI DSC sequence are displayed in **(a)** and **(e)**, respectively. The derived maps of CBV, CBF, and MTT from each sequence are demonstrated subsequently. Note that the DSC CBV and CBF values are not absolute measures and thus are in arbitrary unit (a.u.). The color bars indicate the corresponding scales of each parameter. Note that the spatial resolution in the dual-echo TSE sequence (1×1×2 mm^3^) was higher (about 1/20 in voxel size) than the standard GRE EPI DSC sequence (2.8×2.8×5 mm^3^). The approximately same slice location is shown in the figure.

**Figure 4.**
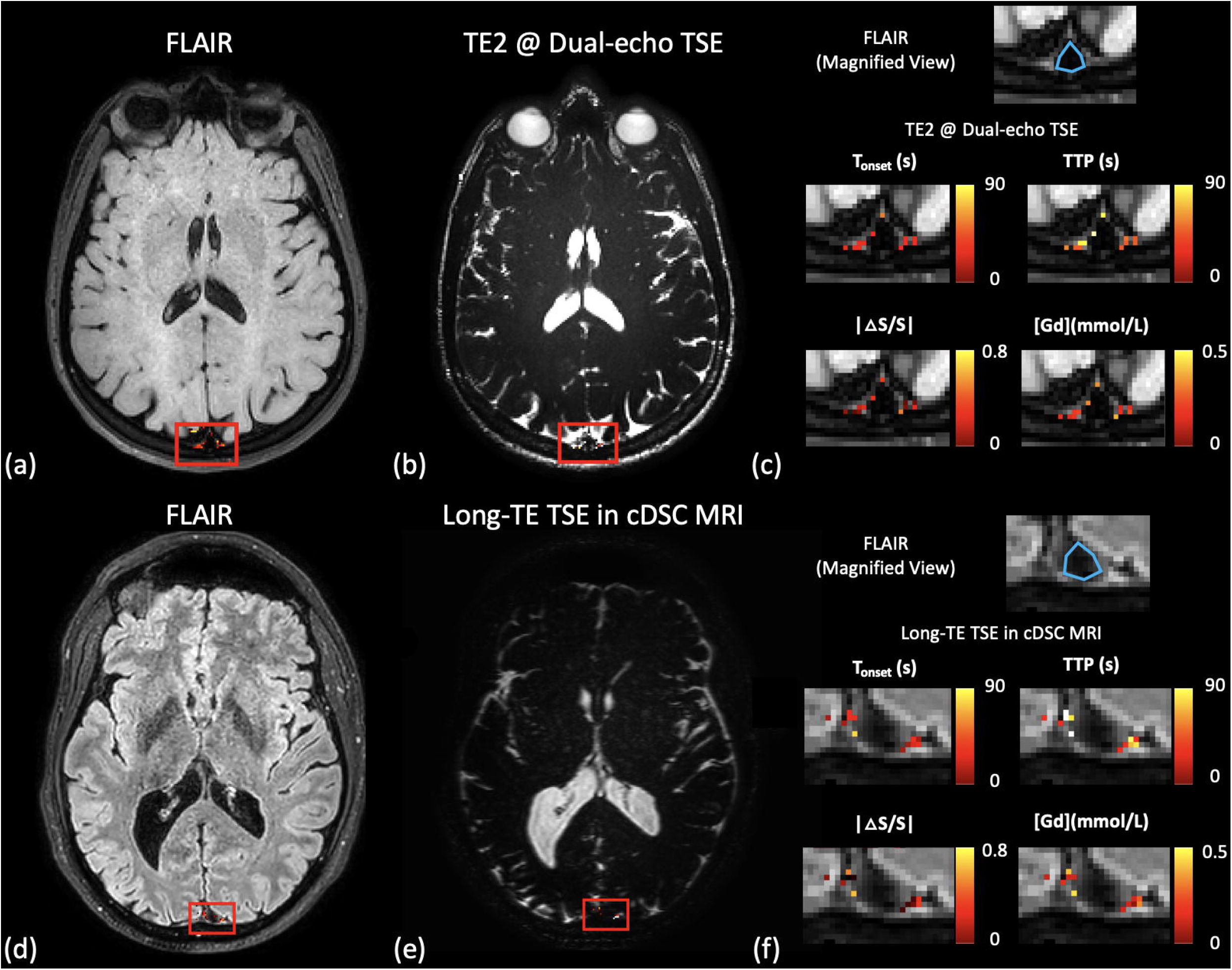
Representative results for the measurement of dynamic signal changes in the CSF from human scans on 3T are shown. The region-of-interest (ROI, red boxes) around the **dural sinuses (DS)** (especially the superior sagittal sinus) that contains the **meningeal** lymphatic vessels is manually drawn. **(a)** The FLAIR image is shown to confirm the location of the meningeal lymphatic vessels. The relative signal changes (ΔS/S) detected with the FLAIR sequence in the ROI are overlaid on the image. **(b)** The raw image acquired at the long TE (TE2 = 560 ms) in the proposed dual-echo TSE sequence is shown with ΔS/S overlaid on the image. Only voxels with a contrast-to-noise ratio (CNR) > 1 were highlighted in the image (see Data analysis). **(c)** A magnified view (axial) of the FLAIR image in the ROI, and maps of the following parameters extracted from the dynamic time courses detected in the dual-echo TSE sequence overlaid on the FLAIR image are shown: T_onset_ = time of onset, TTP = time to peak, absolute value of relative signal change |ΔS/S| between pre- and post-Gd, [Gd] = peak Gd concentration. The blue contour on the magnified FLAIR image outlines approximately the location of the superior sagittal sinus, around which the meningeal lymphatic vessels are located in previous studies. The color bars indicate the corresponding scales of each parameter. For comparison, results from the previous cDSC MRI sequence: the FLAIR image, raw cDSC image, and the corresponding parametric maps are shown in **(d), (e)**, and **(f)**, respectively. The slice location was chosen to cover the same ROI (red boxes) around the **dural sinuses (DS)** (especially the superior sagittal sinus) that contains the **meningeal** lymphatic vessels.

**Figure 5.**
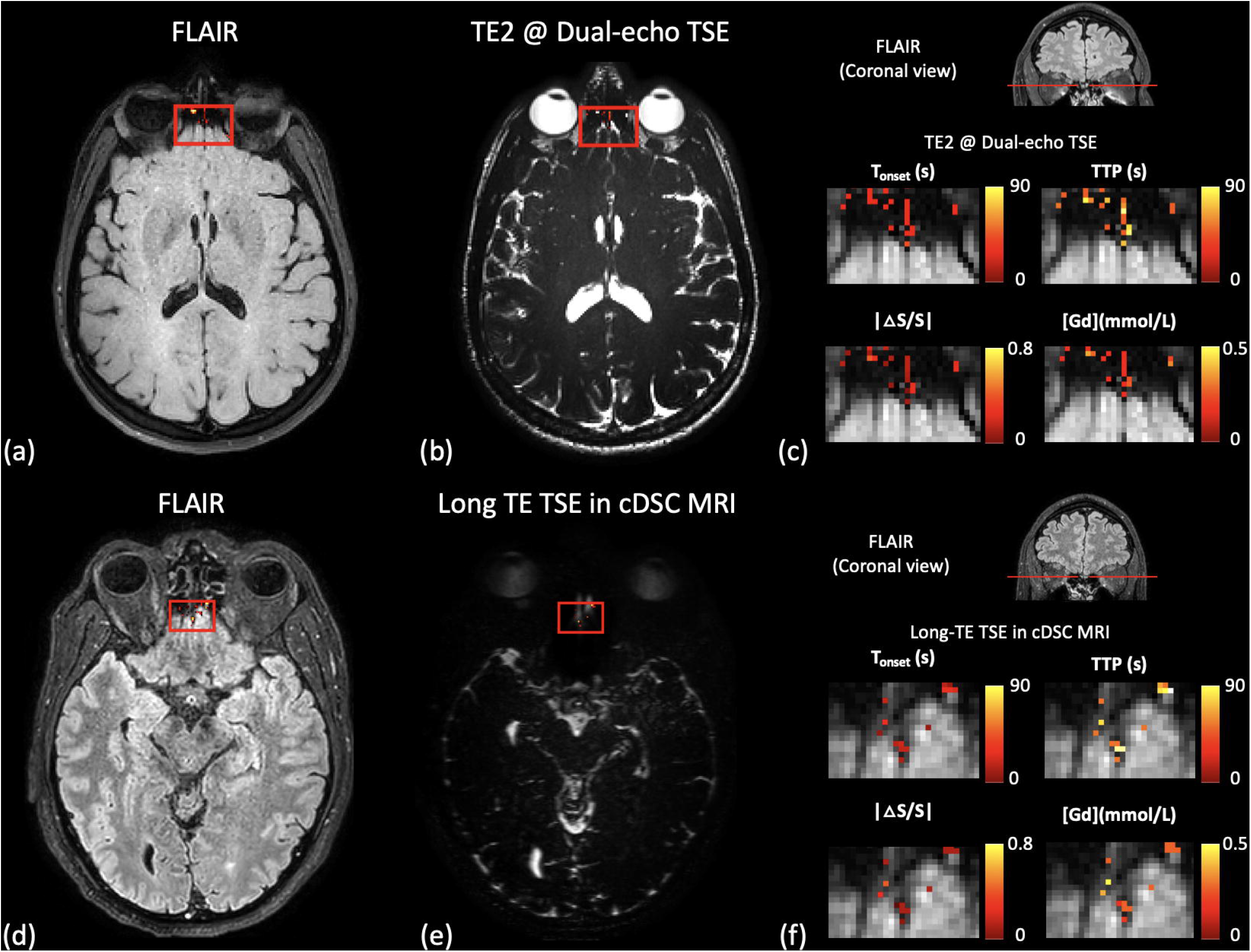
Representative results for the measurement of dynamic signal changes in the CSF from human scans on 3T are shown. The region-of-interest (ROI, red boxes) around the **cribriform plate (CP)** that contains the **olfactory** lymphatic vessels are manually drawn. **(a)** The FLAIR image is shown to confirm the location of the olfactory lymphatic vessels. The relative signal changes (ΔS/S) detected with the FLAIR sequence in the ROI are overlaid on the image. **(b)** The raw image acquired at the long TE (TE2 = 560 ms) in the proposed dual-echo TSE sequence is shown with ΔS/S overlaid on the image. Only voxels with a contrast-to-noise ratio (CNR) > 1 were highlighted in the image (see Data analysis). **(c)** A coronal view of the FLAIR image is shown to indicate the location of the axial slice. Maps of the following parameters extracted from the dynamic time courses detected in the dual-echo TSE sequence overlaid on the magnified axial FLAIR image are shown: T_onset_ = time of onset, TTP = time to peak, absolute value of relative signal change |ΔS/S| between pre- and post-Gd, [Gd] = peak Gd concentration. The color bars indicate the corresponding scales of each parameter. For comparison, results from the previous cDSC MRI sequence: FLAIR image, raw cDSC image, and the corresponding parametric maps are shown in **(d), (e)**, and **(f)**, respectively. The slice location was chosen to cover the same ROI (red boxes) around the **cribriform plate (CP)** that contains the **olfactory** lymphatic vessels.

**Figure 6.**
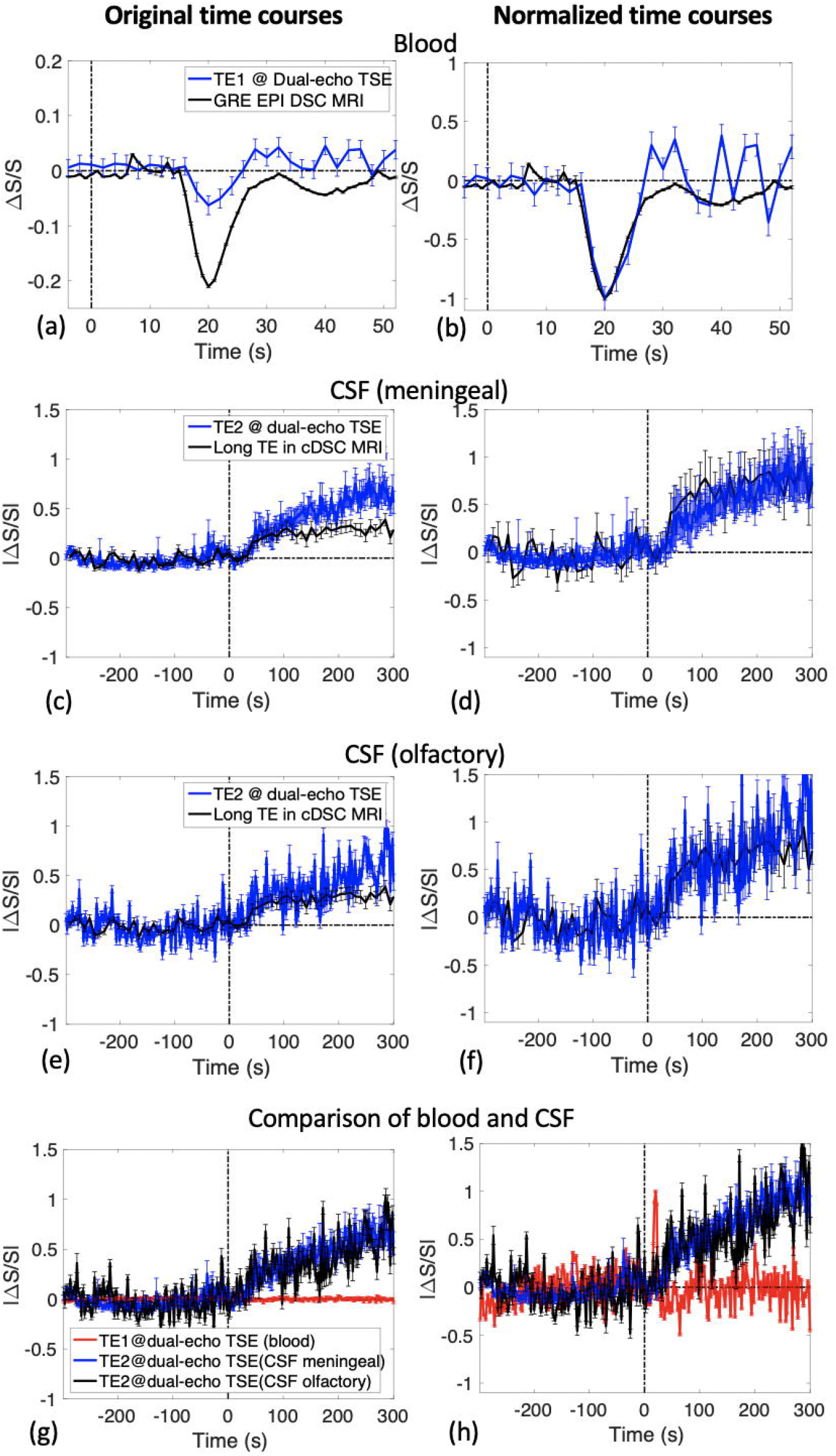
Time courses detected in blood and CSF before and after Gd injection are shown. The relative signal changes (ΔS/S) between pre- and post-Gd images are shown as a function of time. The error bars indicate standard deviations. The vertical dotted lines indicate the time when Gd is injected. **(a)** The average time courses from the blood vessels in grey matter tissue detected using the proposed sequence at short TE (TE1 = 80 ms) and the standard GRE EPI DSC MRI sequence are shown. **(b)** The same time courses as in (a) normalized by their corresponding peaks (the largest signal change) are shown in order to compare their temporal difference. **(c)** The average time courses in the meningeal lymphatic vessels around the dural sinuses region detected using the proposed sequence at long TE (TE2 = 560 ms) and the previous cDSC MRI sequence are shown. **(d)** Normalized time courses from (c). **(e)** The average time courses in the olfactory lymphatic vessels around the cribriform plate detected using the proposed sequence at long TE (TE2 = 560 ms) and the previous cDSC MRI sequence are shown. **(f)** Normalized time courses from (e). **(g)** To compare the time courses in the blood and meningeal and olfactory lymphatic vessels detected using the proposed approach, the same time courses as in (a, c, e) are displayed together. **(h)** Normalized time courses from (g). Note that in **(c-h)**, the absolute values of relative signal changes (ΔS/S) between pre- and post-Gd images are shown for comparison since ΔS/S from different sequences can have opposite signs.

**Table 2.** Quantitative results from DDSEP MRI and the existing separate methods (DSC MRI and cDSC MRI for blood and lymphatic vessels, respectively) in healthy subjects (n = 8).

|  | Separate | Combined | P |
| --- | --- | --- | --- |
| *Cerebrovascular perfusion parameters in grey matter tissue* |  |  |  |
| **tSNR** | 42.7±11.3 | 13.4±7.6 | 0.001* |
| **CNR** | 27.1±3.5 | 5.2±1.3 | < 0.0001* |
| **CBV<sub>GM</sub>/CBV<sub>WM</sub>** | 1.92±0.46 | 1.88±0.69 | 0.91 |
| **CBF<sub>GM</sub>/CBF<sub>WM</sub>** | 2.31±0.31 | 2.26±0.74 | 0.89 |
| **MTT (s)** | 4.73±0.76 | 4.69±0.89 | 0.94 |
| **ΔS/S** | -0.22±0.07 | -0.06±0.03 | 0.0005* |
| **T<sub>onset</sub> (s)** | 15.7±0.8 | 15.6±0.9 | 0.86 |
| **TTP (s)** | 20.4±0.7 | 20.3±0.9 | 0.70 |
| **T<sub>return</sub> (s)** | 28.2±1.0 | 28.6±1.4 | 0.61 |
| *Cerebral lymphatic vessels in the region around the dural sinuses (DS)* |  |  |  |
| **tSNR** | 14.7±4.9 | 13.6±5.7 | 0.75 |
| **CNR** | 2.0±0.4 | 3.1±0.5 | 0.004* |
| **ΔS/S** | -0.28±0.06 | 0.53±0.12 | 0.003* |
| **Peak [Gd] (mmol/L)** | 0.14±0.07 | 0.16±0.06 | 0.62 |
| **T<sub>onset</sub> (s)** | 27.3±8.8 | 29.8±7.4 | 0.67 |
| **TTP (s)** | 52.7±22.4 | 55.4±10.5 | 0.81 |
| *Cerebral lymphatic vessels in the region around the cribriform plate (CP)* |  |  |  |
| **tSNR** | 10.4±5.5 | 11.8±3.2 | 0.64 |
| **CNR** | 1.7±0.5 | 2.3±0.4 | 0.05* |
| **ΔS/S** | -0.33±0.08 | 0.49±0.21 | 0.15 |
| **Peak [Gd] (mmol/L)** | 0.28±0.08 | 0.23±0.09 | 0.36 |
| **T<sub>onset</sub> (s)** | 28.7±23.8 | 32.3±10.3 | 0.60 |
| **TTP (s)** | 71.8±32.9 | 68.9±23.1 | 0.88 |

CBV<sub>GM</sub>/CBV<sub>WM</sub> = cerebral blood volume (CBV) in grey matter (GM) normalized by CBV in white matter (WM); CBF<sub>GM</sub>/CBF<sub>WM</sub> = cerebral blood flow (CBF) in grey matter (GM) normalized by CBF in white matter (WM); MTT = mean transit time; T<sub>onset</sub> = time of onset; TTP = time to peak; T<sub>return</sub> = time of return; tSNR, CNR, relative signal change ΔS/S, and peak Gd concentration ([Gd]) are defined in Methods.

Figure 3. compares DDSEP images acquired at short TE (**Figure 3a**) with the standard GRE EPI DSC images (**Figure 3f**). The parametric maps of CBV, CBF, and MTT for both methods are also compared. Overall, the contrasts in the corresponding images were comparable between the two methods. Note that the spatial resolution in the proposed approach (1×1×2 mm^3^) was much higher (about 1/20 in voxel size) than the DSC MRI scans (2.8×2.8×5 mm^3^, used in a previous study (31)) as such high spatial resolution is preferred for measuring perfusion in lymphatic vessels at the long TE. The time courses averaged over all grey matter voxels (**Figures 6a,b**) showed a consistent temporal pattern from both methods, reaching maximum contrast ~20s after Gd injection and returning to baseline ~28s after Gd injection. **Table 2** compares average tSNR, CNR, relative signal change (ΔS/S) before and after Gd injection, and perfusion parameters related to blood vessels in grey matter measured by the two approaches from all participants. The magnitude of ΔS/S was greater in the standard GRE EPI DSC MRI method mainly due to its gradient echo based contrast as compared to the spin echo based contrast in the proposed TSE sequence. GRE EPI DSC MRI also showed higher tSNR and CNR than the DDSEP sequence (P < 0.05), as is expected from its much larger voxel size (lower spatial resolution) and its gradient echo based contrast. The derived perfusion parameters of CBV, CBF, MTT, T<u>_onset_</u>, TTP, and T_return_ in grey matter were consistent between the two approaches (P > 0.1), and were in the typical range for healthy human subjects as reported in the literature (10).

### Human subject results: dynamic signal changes in the CSF

**Figures 4-5** show typical images acquired using the proposed DDSEP sequence at long TE and the cDSC MRI approach. FLAIR images were used to identify the two ROIs described in Methods, in regions where cerebral lymphatic vessels have been identified in human brains, including around the dural sinuses (superior sagittal sinus) and the cribriform plate (3-5). In both regions, CSF signal changes after Gd injection were detected in FLAIR and in the proposed DDSEP sequences. Maps of several parameters including T_onset_, TTP, ΔS/S, and peak Gd concentration in the CSF ([Gd]) extracted from the dynamic signal changes detected with the DDSEP and cDSC sequences are shown for each ROI. The time courses from both ROIs (**Figures 6c-f**) showed a consistent temporal pattern from both methods, respectively. The CSF signal change after Gd injection detected in both sequences remained preserved for the entire acquisition period (approximately 5 minutes after Gd injection). Note that the ΔS/S in CSF in the DDSEP sequence is expected to be positive (**Figure 1c,d**), whereas ΔS/S in CSF in the original cDSC MRI sequence is expected to be negative (**Figure 1e,f**) due to their T1 and T2 weighting, respectively. Therefore, the absolute values of ΔS/S are compared here. **Table 2** compares average tSNR, CNR and parameters extracted from the dynamic signal changes detected in CSF from the two approaches from all participants. Both approaches showed comparable tSNR (P > 0.1). The magnitude of ΔS/S in CSF was greater in the DDSEP sequence (P < 0.05), which leads to a greater CNR for the proposed method (P < 0.05). The derived temporal parameters of T_onset_ and TTP, and Gd concentration in CSF were consistent between the two approaches (P > 0.1) and in the typical range for healthy human subjects (16).

### Comparison of the dynamic signal changes in blood and lymphatic vessels

The time courses from blood vessels, and meningeal and olfactory lymphatic vessels measured in the proposed DDSEP sequence at the short (TE1) and long (TE2) echoes, respectively, are shown and compared in **Figures 6g,h**. Only time courses measured using the combined method were compared as they are from the same scans and the same subjects. Since ΔS/S in CSF is expected to be positive and ΔS/S in blood negative, absolute values of ΔS/S were compared here. The magnitude of ΔS/S was smaller in blood than in CSF (**Table 2**, P < 0.05). The time of onset (T_onset_) and time to peak (TTP) were both shorter in blood than in CSF (**Table 2**, P < 0.05). The blood time courses quickly returned to baseline whereas the CSF time courses did not return to baseline for the duration of our experiments (approximately 5 minutes after Gd injection). The time of return (T_return_) for blood signals was comparable to the time of onset (T<u>_onset_</u>) for CSF signals (**Table 2**, P > 0.1), and was significantly shorter than the time to peak (TTP) for CSF signals (**Table 2**, P < 0.01).

## Discussion

To date, most existing MRI approaches can only measure perfusion parameters related to blood or lymphatic vessels separately, and most contrast enhanced MRI methods are optimized for detecting Gd-induced signal changes in the blood (10). In the current study, we demonstrate a dual-echo TSE sequence (DDSEP MRI) with optimized contrasts for the detection of Gd induced MR signal changes in the blood and CSF with short (80 ms) and long (560 ms) TE, respectively. A primary advantage of the proposed approach is that it provides a tool for concurrent measurement of perfusion parameters related to both blood and lymphatic vessels in the brain in one single scan, which substantially shortens the total scan time and reduces inter-scan confounding effects from physiological variations. In addition, the dose of contrast medium needed is halved. Importantly, this approach can be used to study the interaction between the microvascular and lymphatic systems in the brain, which plays a critical role in the regulation of blood supply and waste clearance in the brain and has been linked to a number of brain diseases (7-9). Our initial results in healthy human subjects demonstrated that consistent perfusion parameters related to blood and lymphatic vessels can be measured by the proposed combined approach and the respective existing separate methods in the human brain.

Multi-echo based MRI approaches have been employed in DSC and DCE MRI. Dual gradient echo (GRE) based methods have been developed to provide both perfusion (DSC) and vascular permeability (DCE) measures using a single dose of Gd contrast (32-36). The PERfusion with Multiple Echoes and Temporal Enhancement (PERMEATE) method is a dual GRE multi-shot EPI sequence developed to improve R2* quantification and to reduce geometric distortion in typical EPI images (37,38). The Spiral Perfusion Imaging with Consecutive Echoes (SPICE) method is another dual GRE approach using a spiral readout (39). The spin- and gradient-echo (SAGE) sequence combines the acquisition of two GREs, two asymmetric spin echoes, and a single spin echo (40-43) to provide probably a most complete set of hemodynamic parameters related to macro- and microvascular perfusion, vascular permeability, and other properties within a single acquisition (40-44). The simplified SAGE (sSAGE) approach reduces the acquisition to two GREs and a single spin echo, making it more feasible for clinical use (44,45). The key difference in the proposed DDSEP approach is that it is a dual-spin-echo sequence with a long second TE in order to get a pure CSF signal. Thus, the proposed DDSEP sequence furnishes a T1-dominat contrast in the CSF similar to DCE MRI and a T2-dominant contrast in the blood same as DSC MRI. In principle, fast GRE sequences commonly used in standard DCE MRI can provide a T1-dominant CSF contrast as well with some degree of residual T2* effects depending on the TE used. If acquired at multiple gradient echoes, such residual T2* effects can be eliminated by numerically fitting the multi-echo data to an effective TE of 0, thus providing a pure T1 contrast in the CSF. In such fast GRE sequences, both blood and CSF signals are expected to increase after Gd injection. By contrast, in the proposed DDSEP method, the blood and CSF signals show opposite contrasts after Gd injection. Moreover, a main reason for choosing a dual-spin-echo sequence in the proposed DDSEP method that it is known to have better spatial specificity for Gd-induced signal changes in small vessels than GRE-based sequences (27-30), which is critical for the method since cerebral lymphatic vessels often run alongside small blood vessels in the brain (3-5).

The long-TE images in the proposed DDSEP approach provide a clean CSF signal with brain parenchyma and blood signals all suppressed due to T2 decay. Thus, the partial volume effects from brain parenchyma and blood are expected to be minimal in the long-TE CSF images. Therefore, even if the spatial resolution (voxel size) is larger than the dimensions of the target structures (the CSF space and the cerebral lymphatic vessels), dynamic signal changes in the CSF can still be measured with the proposed method. On the other hand, the short-TE images in the proposed DDSEP approach are expected to have a negative and a positive contrast for the blood and CSF signals respectively (see simulations in **Figure 1b**). When the spatial resolution is sufficiently fine to separate the blood and lymphatic vessels, or when imaging brain regions that are expected to have only one type of the microvessels, the short-TE images alone can provide measures of dynamic signal changes in both blood and CSF. However, if the voxel contains both blood and lymphatic vessels, the opposite signal changes in blood and CSF will cancel out with each other, resulting in partial volume effects. As cerebral lymphatic vessels can only be found in several specific brain regions, such partial volume effects in the blood signals in most GM and WM regions are expected to be small. For the CSF signals, the long TE images are required to separate the partial volume effects from blood.

The Gd-induced signal changes in the blood and CSF can be further separated by their differential temporal profiles. According to the data measured in the current study in healthy human subjects (**Figures 6g,h** and **Table 2**), after Gd injection, the blood signals decreased and then quickly restored to baseline before significant changes can be measured in the CSF signals. To the best our knowledge, this may be the first time that dynamic signal changes in small blood and lymphatic vessels in the human brain are measured in the same subjects. The timing parameters measured in the blood and CSF time courses are congruent with the DSC literature (10) and our previous study using cDSC MRI (16), respectively. The timing difference between the blood and CSF signal changes seems to support the hypothesis that after intravenous (IV) injection, the Gd-based contrast agents enter the CSF space via the dural blood vessel wall that lacks a BBB (12-15). This timing difference is also comparable to the timing of signal changes reported in many brain tumor studies using DCE MRI where Gd-based contrast agents often leak out from the impaired BBB in the tumor region (46,47). The exact mechanism for such communication between dural blood vessels and meningeal lymphatic vessels in the parasagittal dura is still under extensive investigation (48). In that regard, the proposed DDSEP approach will be performed around the parasagittal dura region in subsequent studies in order to examine the pathway for Gd-based contrast agents to travel from dural blood vessels to meningeal lymphatic vessels in the human brain.

The timing difference between the blood and CSF signal changes also enables us to estimate Gd concentration ([Gd]) in the CSF using the proposed DDSEP MRI approach. In the original cDSC MRI, Gd concentration can be uniquely determined from the T2-dominant CSF contrast (16). However, the T1-dominant contrast for CSF from the long-TE images in DDSEP MRI has a bi-phasic relationship with Gd concentration (**Figure 2b**), which makes it uncertain to estimate [Gd] from the long-TE images alone. Signal changes from the short-TE images in DDSEP MRI have a monotonic relationship with Gd concentration in the CSF (**Figure 2**) when the Gd concentration is within the typical range expected in the human brain after intravenous (IV) injection of a standard dose of Gd-based contrast agents. With the significant timing difference between the blood and CSF signal changes detected in our data, the blood signal changes can be assumed to be 0 when the CSF signal changes reach the peak. This temporal separation allows us to determine Gd concentration in the CSF from the short-TE images acquired around the CSF TTP in DDSEP MRI. In our data, the peak Gd concentration in CSF estimated using the new DDSEP MRI approach was in agreement with the results from our previous study using the cDSC MRI method (16). One limitation is that if the Gd concentration exceeds the expected range of 0-0.5mmol/L, which is unlikely in human studies but possible in animal studies with different Gd injection methods, or if the Gd-induced blood signal changes concur or overlap with CSF signal changes, additional information is needed for the estimation of Gd concentration in DDSEP MRI.

### Technical limitation and future direction

As a first proof-of-concept study, the proposed DDSEP MRI approach was implemented in multi-slice acquisition mode with 3 slices covering the superior sagittal sinus and the olfactory regions where cerebral lymphatic vessels have been identified. Dynamic signal changes from both blood and lymphatic vessels were detected using DDSEP MRI, based on which we can further optimize and develop the DDSEP MRI sequence in subsequent studies. Although cerebral lymphatic vessels are only identified in several specific brain regions, a whole brain coverage would still be ideal for DDSEP MRI. The main limiting factor here is the long TE (> 500 ms) required for a clean CSF signal and a short TR (< 2 s) required for tracking the dynamic phase of the blood signal. With the multi-slice acquisition mode, only 3 slices can be fit into a TR of 2 s in the current study. A few approaches can be employed to improve the spatial coverage in future studies. First, the flip angles of the readout RF pulse train in the TSE sequence can be optimized to make the required long TE shorter for a complete suppression of blood signals. Second, a three-dimension (3D) acquisition mode with lower spatial resolution can be used instead of the current multi-slice mode. The 3D mode will allow more acceleration using advanced fast imaging techniques such as the Compressed-Sensing SENSE (CS-SENSE) technique (49,50) as in our previous study (16) to improve the spatial coverage with the same readout time. In DDSEP MRI, the short-TE images are mainly used to measure blood perfusion similar to DSC MRI, which is typically performed at a much lower spatial resolution (2-3mm instead of 1mm currently). The spatial resolution for the long-TE images in DDSEP MRI can also be reduced substantially because negligible partial volume effects are expected with blood and parenchymal signals suppressed. Finally, based on our data in healthy human subjects, the dynamic phases for blood and CSF signal changes after intravenous (IV) Gd administration have a significant temporal separation. As the short-TE images can detect Gd-induced signal changes in both blood and CSF as shown in the simulations (**Figure 1b**), such temporal separation seems to eliminate the need for the long-TE images in DDSEP MRI, which will significantly shorten the acquisition time. Furthermore, different readout sequences can be optimized for the blood and CSF phases, respectively. All these options discussed above will be evaluated in our follow-up study.

In addition to CBV and CBF, measures related to the BBB permeability are quite useful in various brain diseases. In the current DDSEP MRI sequence, the short TE is chosen to produce T2-weighted images for the blood signals. Therefore, the K2 parameter from DSC MRI can be calculated using the short-TE images provided sufficient SNR is achieved. However, the more commonly used K_trans_ parameter for permeability from DCE MRI would need a T1-weighted blood contrast. To obtain such T1-weighted blood contrast, images need to be acquired at a TE much shorter than the current TE1 (= 80 ms). Alternatively, one can also acquire images at an additional TE in the intermediate range (100-300 ms), and numerically fit for an effective TE of 0 using data from both echoes. The long TE in the current DDSEP MRI sequence is too long for blood signals which have already decayed to almost zero, and thus cannot be used for multi-echo fitting. These technical development is being pursued in follow-up studies.

In summary, we demonstrated the DDSEP MRI approach based on a dual-echo TSE sequence which can measure perfusion parameters related to both small blood and lymphatic vessels concurrently after intravenous (IV) injection of Gd contrast medium in healthy human subjects. The results from the DDSEP MRI method are in line with previous studies using separate methods for small blood and lymphatic vessels, respectively. Interestingly, signal changes from small blood vessels occurred much faster than those from the CSF and cerebral lymphatic vessels. To the best of our knowledge, this may be the first study in which such interaction was measured and reported in the same human subjects. The proposed methodology is being further developed to expand its spatial coverage to the whole brain. Given the importance of the microvascular and cerebral lymphatic systems in the brain and their tight interaction, we believe that the proposed DDSEP MRI approach may provide a useful tool for studies on these systems in the healthy brain and various brain diseases.

## Acknowledgments

The authors thank Mr. Joseph S. Gillen, Mrs. Terri Lee Brawner, Ms. Kathleen A. Kahl, and Ms. Ivana Kusevic for experimental assistance. This project was supported by the National Institutes of Health through grants from the NINDS (1R01NS108452 and 1R01NS120879), NIA (5R01AG064093), the NIBIB (P41 EB015909), and the NICHD (U54 HD079123). Equipment used in the study was manufactured by Philips. Under a license agreement between Philips and the Johns Hopkins University, Dr. Knutsson’s spouse, Dr. van Zijl and the University are entitled to fees related to an imaging device used in the study discussed in this publication. Dr. van Zijl also is a paid lecturer for Philips. This arrangement has been reviewed and approved by the Johns Hopkins University in accordance with its conflict of interest policies.

