## Appendix for "Concurrent measurement of perfusion parameters related to small blood and lymphatic vessels in the human brain using dynamic dual-spin-echo perfusion (DDSEP) MRI"

**Supporting Information:**

**MR signal equations in the proposed dual-echo TSE MRI sequence.**

The MR signal (S) measured in a TSE sequence is related to the magnetization (M_z_) with the following equation:

$S\sim M_{z}=M_{0}\times(1-e^{-\frac{TR}{T1}})\times e^{-\frac{TE}{T2}}$	[S1]

in which M_0_ is the equilibrium longitudinal magnetizations. Note that TE here represents the effective echo time which is determined by the readout echo train in the sequence (1).

The Gd induced changes in T1 and T2 relaxation times can be written as:

$\frac{1}{{T1}_{Gd}}= \frac{1}{{T1}_{0}}+r_{1}\times[Gd]$	[S2]

$\frac{1}{{T2}_{Gd}}= \frac{1}{{T2}_{0}}+r_{2}\times[Gd]$	[S3]

where variables with the subscription “0” (T1_0_ and T2_0_) represent the values before Gd injection, variables with the subscription “Gd” (T1_Gd_ and T2_Gd_) represent the values after Gd injection, r1 and r2 are the relaxivity of Gd in a certain medium (CSF or blood), and [Gd] is the concentration of Gd in the medium.

In the proposed dual-echo TSE sequence, the MR signal measure at the short TE (TE1) can be written as:

Before Gd injection:

${S1}_{0}\sim X_{blood}\times C_{blood}\times M_{blood}+X_{CSF}\times C_{CSF}\times M_{CSF}+\left( {1-X}_{blood}-X_{CSF} \right)\times C_{tissue}\times M_{tissue}$	[S4a]

After Gd injection:

${S1}_{Gd}\sim X_{blood}\times C_{blood}\times M_{blood,Gd}+X_{CSF}\times C_{CSF}\times M_{CSF,Gd}+\left( {1-X}_{blood}-X_{CSF} \right)\times C_{tissue}\times M_{tissue}$	[S4b]

where Xi (i = blood, CSF) is the compartmental fraction (ranging from 0 to 1), Ci (i = blood, CSF, tissue) is the water density for each compartment, and Mi (i = blood, CSF, tissue) is defined in Eq. [S1] with respective T1 and T2 values.

At the long TE (TE2) in the proposed dual-echo TSE sequence, MR signals from blood and tissue can be assumed zero due to T2 delay. Therefore, the equations can be simplified to:

${S2}_{0}\sim X_{CSF}\times C_{CSF}\times M_{CSF}=X_{CSF}\times C_{CSF}\times M_{0}\times(1-e^{-\frac{TR}{T1,CSF,0}})\times e^{-\frac{TE2}{T2,CSF,0}}$	[S4c]

${S2}_{Gd}\sim X_{CSF}\times C_{CSF}\times M_{CSF,Gd}=X_{CSF}\times C_{CSF}\times M_{0}\times(1-e^{-\frac{TR}{T1,CSF,Gd}})\times e^{-\frac{TE2}{T2,CSF,Gd}}$	[S4d]

There are two unknown variables in Eqs. [S4c,d]: X_CSF_ and [Gd] (which gives rise to T_1,CSF,Gd_ and T_2,CSF,Gd_). Therefore, in principle, with the two measures S2_0_ and S2_Gd_, X_CSF_ and [Gd] can be obtained from Eqs. [S4c,d]. However, because S2_Gd_ has a biphasic relationship with [Gd] (see simulation results presented in **Figure 2b**), [Gd] cannot be uniquely determined with S2_Gd_ alone.

On the other hand, within the range of [Gd] (0-0.5 mmol/L) expected in the CSF and cerebral lymphatic vessels in the human brain after intravenous (IV) Gd administration with a standard dose, S1_Gd_ increases with [Gd] monotonically (see simulation results presented in **Figure 2b**). Furthermore, as shown in our data from healthy human subjects in **Figure 6**, the blood and CSF signal changes showed significant temporal separation for at least 20 seconds. When the CSF signals showed significant Gd-induced changes, the blood signals have returned to its pre-Gd level. Therefore, M_blood_ in S1_0_ and M_blood,Gd_ in S1_Gd_ can be assumed the same 30 seconds after Gd injection. Based on these assumptions, the absolute signal change (ΔS) before and after Gd injection measured at TE1 can be defined as:

$\Delta S1={S1}_{Gd}-{S1}_{0}\sim X_{CSF}\times C_{CSF}\times\left( M_{CSF,Gd}-M_{CSF} \right)=X_{CSF}\times C_{CSF}\times M_{0}\times\left[ \left( 1-e^{-\frac{TR}{{T1}_{CSF,Gd}}} \right)\times e^{-\frac{TE1}{{T2}_{CSF,Gd}}}-\left( 1-e^{-\frac{TR}{{T1}_{CSF,0}}} \right)\times e^{-\frac{TE1}{{T2}_{CSF,0}}} \right]$	[S5]

Because ΔS1 has a monotonic relationship with [Gd] (**Figure 2b**), [Gd] can be uniquely solved from ΔS1/S2_0_:

$\frac{\Delta S1}{{S2}_{0}}=\frac{\left( 1-e^{-\frac{TR}{{T1}_{CSF,Gd}}} \right)\times e^{-\frac{TE1}{{T2}_{CSF,Gd}}}-\left( 1-e^{-\frac{TR}{{T1}_{CSF,0}}} \right)\times e^{-\frac{TE1}{{T2}_{CSF,0}}}}{(1-e^{-\frac{TR}{T1,CSF,0}})\times e^{-\frac{TE2}{T2,CSF,0}}}$	[S6]

Note that S2_0_ can be obtained from the long TE (TE2) in the proposed dual-echo TSE sequence, or it can simply be acquired using a separate scan with the same parameters before Gd injection.
